# SIRT1 activity orchestrates ECM expression during hESC-chondrogenic differentiation

**DOI:** 10.1101/2020.05.12.087957

**Authors:** Christopher A Smith, Paul A Humphreys, Nicola Bates, Mark A Naven, Stuart A Cain, Mona Dvir-Ginzberg, Susan J Kimber

## Abstract

Epigenetic modification is a key driver of differentiation and the deacetylase Sirtuin1 (SIRT1) is an established regulator of cell function, ageing and articular cartilage homeostasis. Here we investigate the role of SIRT1 during development of chondrocytes by using human embryonic stem cells (hESCs). HESC-chondroprogenitors were treated with SIRT1 activator; SRT1720, or inhibitor; EX527, at different development stages. Activation of SIRT1 during 3D-pellet culture led to significant increases in expression of ECM genes for type-II collagen (*COL2A1*) and aggrecan (*ACAN*), and chondrogenic transcription factors *SOX5* and *ARID5B*, with SOX5 ChIP analysis demonstrating enrichment on the ACAN –10 enhancer. Unexpectedly, while ACAN was enhanced, GAG retention in the matrix was reduced when SIRT1 was activated. Significantly, *ARID5B* and *COL2A1* were positively correlated, with Co-IP indicating association of ARID5B with SIRT1 suggesting that COL2A1 expression is promoted by an ARID5B and SIRT1 interaction. In conclusion, SIRT1 activation positively impacts on the expression of the main ECM proteins, whilst altering ECM composition and suppressing GAG content during cartilage development. These results suggest that SIRT1 activity can be beneficial to cartilage development and matrix protein synthesis but tailored by addition of other positive GAG mediators.

## Introduction

Human embryonic stem cells (hESCs) and induced pluripotent stem cells (iPSCs), together termed human pluripotent stem cells (hPSC), have great potential for understanding human development including of skeletal tissues, because of their self-renewal properties and ability to differentiate into many different tissue lineages [1,2]. We have previous reported the differentiation of chondroprogenitors and chondrocytes from hESCs [3] based on developmental principles [4,5]. The cells generated express the key chondrogenic transcription factors SOX 9, 5 and 6 as well as the archetypal matrix component type-II collagen. Additionally, in keeping with a hyaline-like differentiation and the generation of articular cartilage, type X collagen (COLX) is undetectable, indicating that hypertrophy and growth plate like cartilage is not being generated. Whilst promising, expression of the hyaline cartilage proteoglycan aggrecan in the matrix remains low suggesting incomplete matrix synthesis and immaturity so it is important to remedy this to produce a valid developmental model.

Epigenetic effectors can regulate gene expression by modifying histones, as well as various transcription factors. Therefore, modulation of these epigenetic effectors may influence gene expression in chondroprogenitors. The NAD dependent deacetylase, SIRT1 is an epigenetic effector capable of deacetylating histones, and non-histone proteins, such as P53 [6] and RelA/NF-◻B [7], the latter implicated in loss of pluripotency [8]. Importantly, the major transcription factor driving cartilage formation, SOX9 [9,10] is a target of SIRT1 [11,12], which deacetylates it, promoting chondrogenic activity [13].

While much is known about the role of SIRT1 in cartilage homeostasis and disease, much less is known about its role in development. SIRT1 expression is required for the maintenance of hyaline cartilage, and chondrocyte differentiation [12,13] and has been identified as a pro-survival and - metabolic factor, maintaining the homeostasis of the adult chondrocyte in its niche [14]. Importantly, *Sirt1*^−/−^ knock out mice display skeletal and cartilage matrix deficiencies [15–17], indicating a positive role of SIRT1 in articular cartilage homeostasis, but also an involvement in the initial cartilage ECM expression during development. SIRT1 may achieve this by repressing catabolic gene expression [14] while promoting anabolic pathways towards collagen type-II and aggrecan. However, there is a lack of knowledge of the timing and role of SIRT1 in human cartilage development. Using RNAseq analysis of hESC-chondroprogenitors, we identified the presence of SIRT1 alongside significant increases in supportive chondrogenic transcription factors, SOX5 and AT-Rich Interaction Domain 5B (ARID5B), suggesting their importance [18]. SOX5 and ARID5B are both able to form complexes with SOX9 to facilitate chondrocyte maturity. Specifically, SOX9 participates with SOX5 in the SOX trio [19–21] to direct aggrecan expression, while ARID5B binds the epigenetic factor (PHF2) to direct aggrecan and type-II collagen expression [22]. Here we assess the role of SIRT1 in regulating aggrecan and collagen type-II expression during hESC chondrogenesis. Specifically, by modulating SIRT1 activity during the chondrogenesis protocol we start to decipher the pathway by which it regulates these key chondrogenic genes and its role in human cartilage development.

## Results

### SIRT1 expression is dynamic, and is elevated during early stages of chondrogenic differentiation

To understand the importance of SIRT1 in early chondrogenic development, we first sought to determine its expression during the generation of chondroprogenitors, using the 14-day developmental protocol previously published (DDP). and shown in Fig 1. In this protocol cells are induced, by serial growth factor application, through a series of developmental steps to a primitive streak like stage, then to mesodermal progenitor followed by chondroprogenitors [3–5,18].

**Figure 1.**
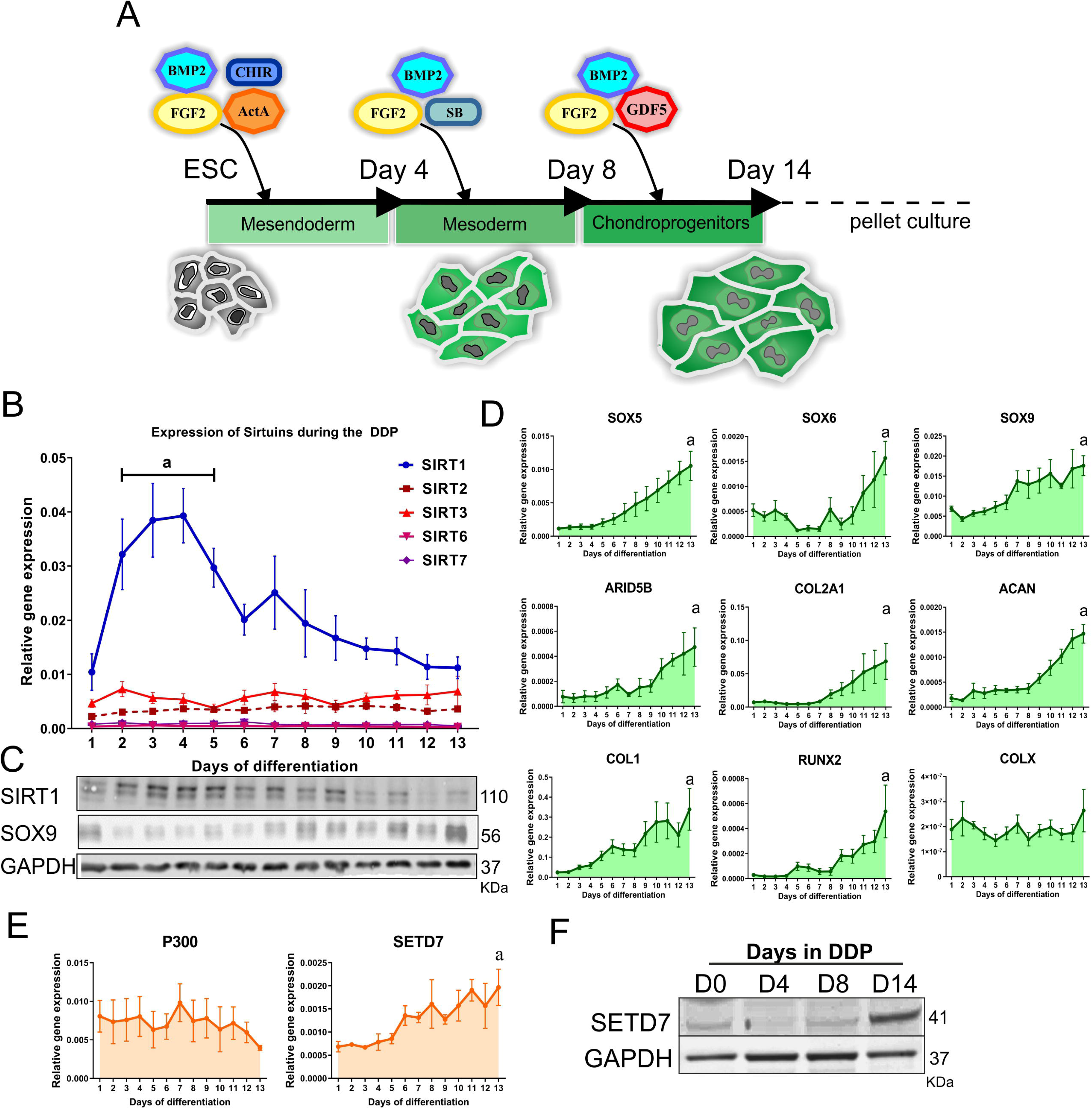
SIRT1 expression changes during the chondrogenic differentiation of hPSCs to chondroprogenitors. A) Schematic diagram of 2D phase of DDP differentiation protocol B) QRT-PCR gene expression analysis of Sirtuins 1,2,3,6 and 7 in samples taken daily (D1-13) over the 13 days of the DDP chondrogenic differentiation process (N=4 biological repeat). C) Western blot analysis of protein expression for SIRT1, SOX9, and the housekeeping control gene GAPDH in samples taken daily from the chondrogenic DDP protocol. Samples correspond to days displayed in panel (B) above. D) QRT-PCR gene expression analysis of transcripts associated with formation of permanent cartilage (*SOX5*, *SOX6*, *SOX9*, *ARID5B*, *COL2A1* and *ACAN*) and other fibrocartilage associated phenotypes (*COL1*, *RUNX2*, *COLX*) in samples taken daily form the DDP chondrogenic protocol (N=4 biological repeat). E) QRT-PCR gene expression and western blot (F) for protein expression of epigenetic factors associated with chondrogenic regulatory mechanisms in chondrogenesis. Data displayed as relative gene expression to housekeeping gene *GAPDH* and shown in box plot. *a* indicates significant difference to day 0 (p≤0.05).

SIRT1 gene and protein expression significantly increased between days 2-5 of differentiation, then decreased steadily back to the original level by day 14 (Figure 1B, C). Alternate Sirtuins were also assessed, however their transcript expression did not significantly change during the differentiation process (Figure 1B). In agreement with our previous studies [5], transcript expression for several chondrogenic genes increased steadily throughout the protocol (Figure 1D) including the core ECM components *COL2A1* and *ACAN*, *ARID5B,* and the major chondrogenic transcription factors *SOX5, SOX6* and *SOX9*. In parallel, we also observed an increase in SOX9 protein levels throughout the protocol (Figure 1C). The hypertrophic marker COLX was not detected during the differentiation process.

Since P300 and SETD7 are associated with epigenetic regulation in chondrogenesis [11,23], and both directly interact with SIRT1 on the COL2A1 promoter [24,25], these were also evaluated. A significant increase in SETD7 by day 14 of the protocol was observed, whilst no change in *P300* gene expression was reported as shown in Figure 1E and 1F. Moreover, we observed no significant change in the gene expression of other chondrogenic-associated enhancing (*HDAC2, HDAC4, TIP60* and *KDM4B*), or repressing (DNA methyltransferase 1 (*DMNT1), KDM2B*) epigenetic modifiers (Figure S1A). Whilst transcript and protein levels of the ARID5B cofactor; histone demethylase PHF2 [22,26] were also detectable at day 14 of differentiation (Figure S1A and B). These data confirm that the increase in chondrogenic genes during differentiation is paralleled by changes in a subset of associated epigenetic effectors.

### SIRT1 inhibition correlates with elevated *SOX9* transcription in hPSC chondrogenesis

Next, we assessed the influence and timing of early SIRT1 activity on eventual chondrogenic differentiation by modulating SIRT1 activity during days 2-5 (stage1), or late from day 8 onwards of the monolayer (2D) protocol (stage 3). Following the administration of the selective SIRT1 inhibitor EX527 [13,27] during stage 1, *SOX9* gene expression increased slightly at day 14 compared to same day control, however SOX9 protein level remained unchanged (Figure S1C, D). Inhibition of SIRT1 during stage 3 (i.e. from day 8-14) had no effect on expression levels of evaluated target genes at day 14 (Figure S1E). Activation of SIRT1 by the highly specific activator SRT1720 [28,29] at either days 2-5 or from day 8-14 in all groups(stage 3) did not results in any significant difference to control (Figure S1C, E). Together, the results of SIRT1 modulation during the early chondrogenic 2D differentiation protocol, support the idea that transition from mesoderm to a chondroprogenitor, during stages 1 and 2 of the protocol, does not require SIRT1 activity, indeed, potentially it could be detrimental since its inhibition led to an increase in expression of the chondrogenic transcription factor SOX9.

### Activation of SIRT1 in a three-dimensional (3D) pellet culture increased chondrogenic gene expression

As previous reports showed that SIRT1 modulation in a 3D setting resulted in augmented COL2 expression [23], we next took the D14 hESC-chondroprogenitors and assessed them in a 3D culture system. Specifically, after 14 days of differentiation (Figure 1A) hESC-chondroprogenitors were pelleted, cultured for 3 days to establish 3D pellets, then the pellets were incubated in chondrogenic culture medium containing SIRT1 activity modulators SRT1720 or EX527 from then on (Figure 2A). Results indicated that SIRT1 activation using 5 μM SRT1720, increased chondrogenic gene expression compared to untreated controls during the 7 days of pellet culture (Figure S2A).

**Figure 2.**
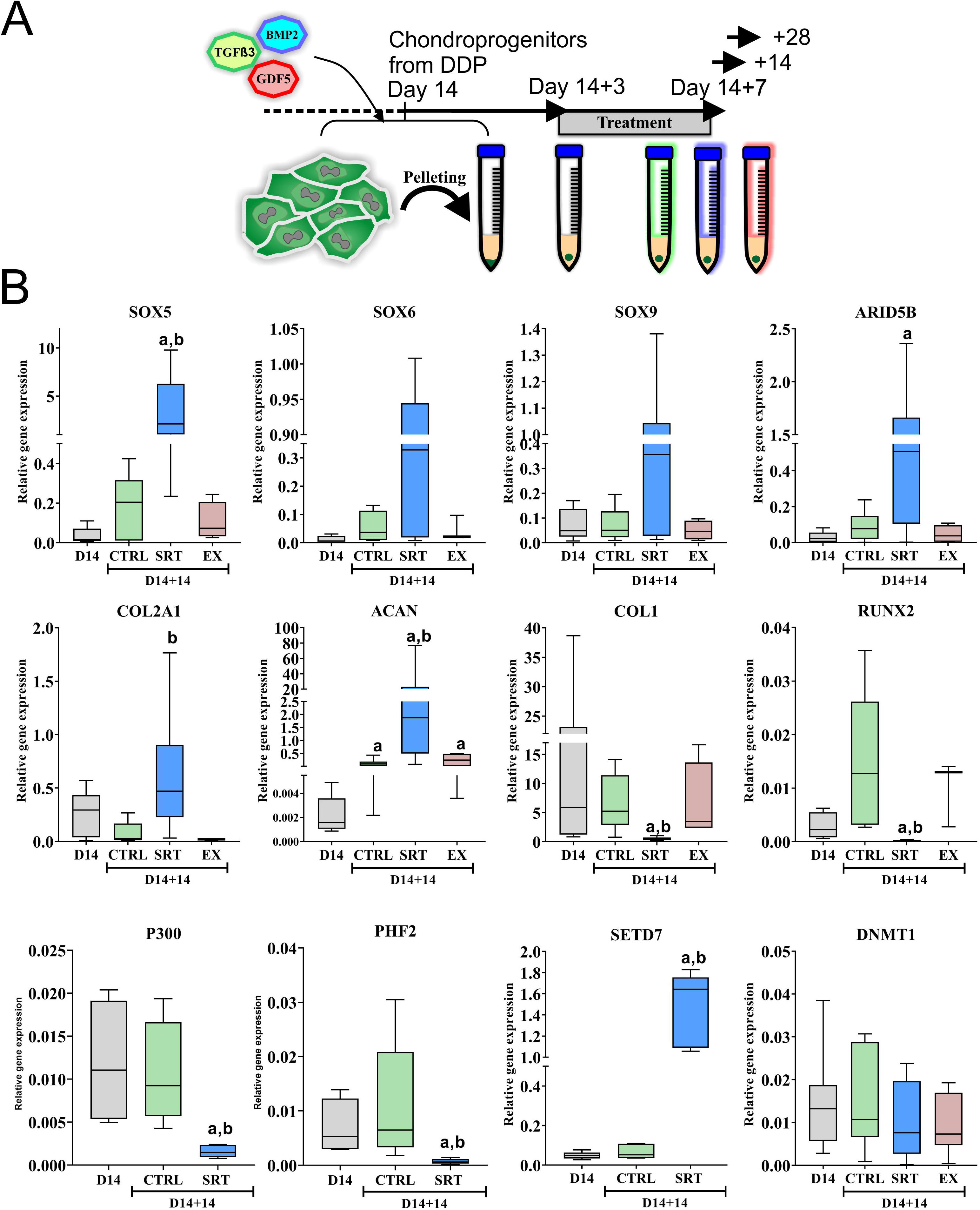
SIRT1 activation in hPSC derived pre-chondrocyte pellets modulates chondrogenic ECM expression. A) Schematic of pellet formation and 3D culture phase of DDP differentiation protocol. B) QRT-PCR gene expression analysis of chondrogenic and non-chondrogenic associated genes in 2D day 14 pre-pellet cells, and pellets at day 14+14, treated with DMSO control or SIRT1 activity modulators from day 14+3 to 14+14 (N=7 biological repeat). Data displayed as gene expression relative to housekeeping gene *GAPDH* and shown in box plot. *a* indicates significant difference to Day 14 pre-pellet sample (p≤0.05), *b* indicates significant difference to day14+14 DMSO vehicle control (p≤0.05).

Gene expression of pellets treated between 3 and 14 days after pelleting revealed a significant increase in expression of chondrogenic genes (i.e. *SOX5*, *ARID5B*, *COL2A1* and *ACAN*) in SRT1720 treated pellets compared to the starting chondroprogenitors (day 14 monolayer (2D) cultures), and untreated pellet controls. Conversely, there was a significant decrease in expression of the fibroblast/hypertrophy-associated genes *COL1* and *RUNX2* in these pellets compared to untreated pellet controls (Figure 2B), while *COLX* was not detected in any of these conditions (data not shown). Notably, these changes were not evident when pellets were treated with the activator for a shorter period of 3 days later on (Figure S2B).

Interestingly, gene expression for both *P300* and *PHF2* associated with SIRT1 action was significantly decreased after SRT1720 treatment compared to that in pre-pelleted cells or untreated pellet controls, whilst an independent epigenetic regulator, DMNT1, which is not associated with SIRT1, was unaffected (Figure 2B). In contrast, *SETD7* expression was significantly increased after SIRT1 activation. However inhibiting SETD7 methyl-transferase activity with (r)-PFI-2 in pellet culture did not affect chondrogenic gene expression compared to control (Figure S2C), indicating that SETD7 activity is likely not essential to SIRT1 action at this stage. Together, this data indicates that SIRT1 selectively regulates other epigenetic modulators as well as recognized chondrogenic genes.

### SIRT1 activation alters cartilage ECM expression and pellet histology

Prolonged SIRT1 activation (i.e. between 3 and 14 days after pelleting) resulted in enlarged pellets compared to controls, while pellets treated with EX527 did not display any change in size (Figure 3A). Histological analysis of control pellets displayed lacunae type structures, which were abundant throughout the pellet, while elongated cells were observed on the pellet surface, reminiscent of an articular surface (Figure 3B). Activation of SIRT1 reduced the abundance of these lacunae in the pellets and decreased alcian blue staining and GAG content (Figure S3A). compared to control pellets, despite a clear increase in antibody staining for aggrecan (Figure 3B, enlarged view 3C). Similarly, hyaluronic acid HA and the major enzyme driving HA synthesis, HAS2, protein were reduced after SIRT1 activation (Figure S3B-C). As well as the increase in type-II collagen, aggrecan and lubricin expression in pellets (Figure S3C), there was also an increase in cells with nuclear staining for SOX5 in SRT1720 treated pellets (Figure 3B), correlating with its elevated gene expression (Figure 2B). This indicates a differential regulation by SIRT1 between chondrogenic proteins and GAGs during hESC-chondrocyte differentiation.

**Figure 3.**
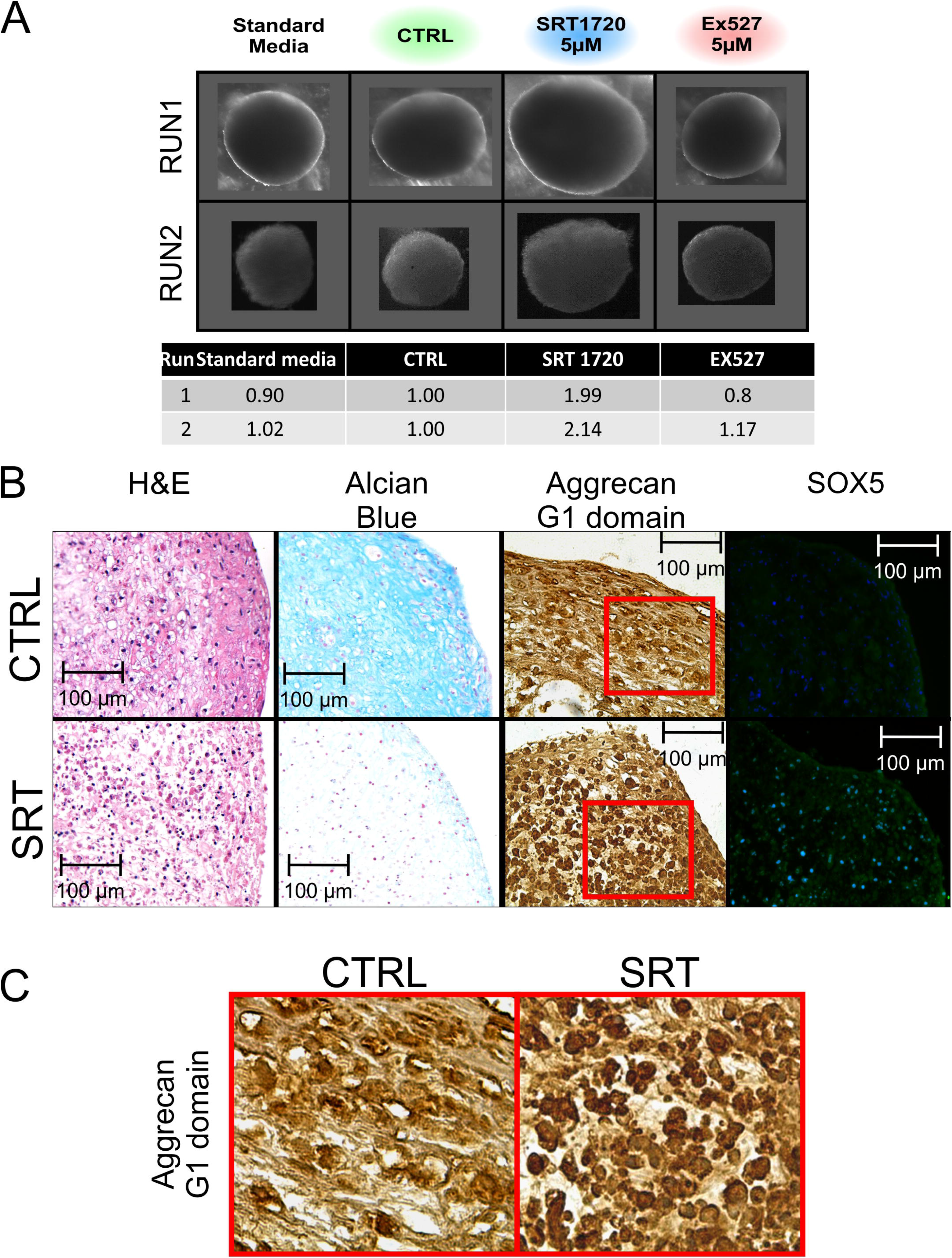
SIRT1 activation changes pellet structure and ECM component aggrecan expression. A) Images of 3D pellet cultures at day 14+14 treated with vehicle controls or SIRT1 activity modulators; SRT1720 SIRT1 activator, or EX527 SIRT1 inhibitor, and table depicting volume relative to standard medium control, from day 14+3 to 14+14. B) IHC and pellet histology for structure (H&E) and GAG staining (alcian blue), immunohistochemical staining for Aggrecan G1 domain and immunofluorescence SOX5 in day 14+28 pellets in 3D cultures treated +/− SRT1720 (5 μM). C) Enlarged view of aggrecan G1 domain staining. Position indicated by red outline.

### Activation of SIRT1 in TC28a2 3D pellets enhances chondrogenic gene expression

Given that SIRT1 activation of hESC-chondroprogenitors induced chondrogenic gene expression in 3D pellets, we investigated whether this response was dependent on developmental maturity using a mature chondrogenic cell line (TC28a2). Monolayer or pelleted TC28a2 cells were cultured for 3 days, before treatment with SRT1720 or vehicle control for an additional 4 days. Results indicated a small but insignificant increase in chondrogenic gene expression in TC28a2 pellets compared to monolayer cultures (Figure 4A). SIRT1 activation in TC28a2 pellets induced a significant increase in transcript for chondrogenic genes compared to the control (Figure 4A), whilst there was no observable change in gene expression in SRT1720 treated monolayer cells. *COL1A2* expression was not affected by SIRT1720 in either culture format. Assessment of aggrecan protein levels by western blotting indicated a substantial increase in the band at 80KDa [30], in line with the increased level of *ACAN* gene expression (Figure 4B). Hence, these results show that SIRT1 activation increased ACAN gene expression during chondrogenesis in a 3D setting, similar to in hESC-chondroprogenitor pellets.

**Figure 4.**
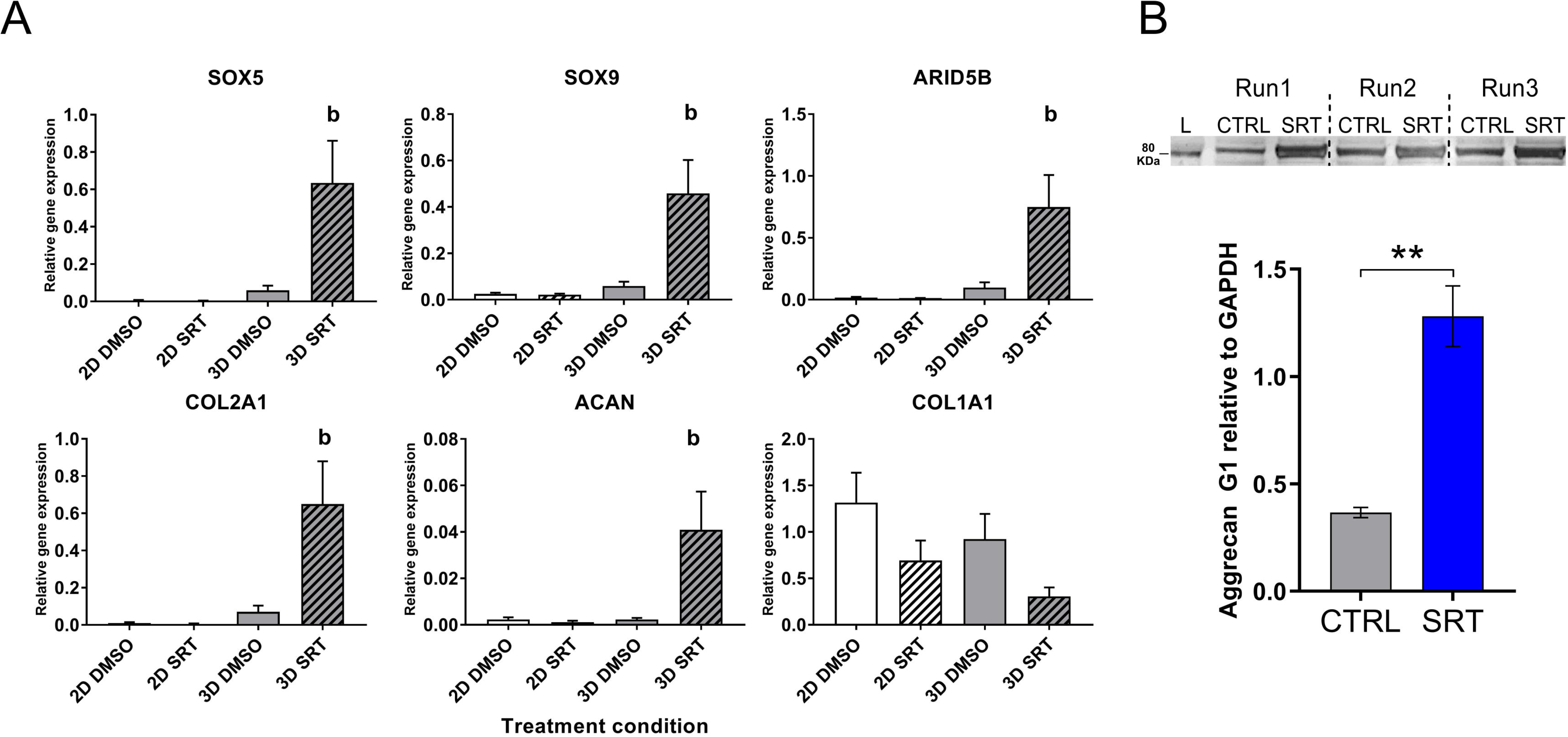
Activation of SIRT1 in TC28a2 cells enhances chondrogenic gene expression in 3D but not 2D culture. A) QRT-PCR gene expression analysis of chondrogenic associated genes in TC28a2 immortalised chondrocytes in 2D or 3D culture for 7 days, treated with DMSO vehicle control or SIRT1 activator SRT1720 (5 μM) from days 3-7 (N=3 biological repeat). B) Western blot protein expression analysis of G1 domain of aggrecan in TC28a2 14-day old pellets treated with DMSO or SRT1720 from day 3 till 14, and quantification of densitometry (N=3 biological repeat). Gene expression data displayed as relative to housekeeping gene *GAPDH. b* indicates significant difference to 3D DMSO vehicle control (p≤0.05). ** indicates significant difference (p≤0.01) compared to control.

### SIRT1 activity not expression level drives chondrogenic gene expression

Based on the above results, we assessed whether overexpression of SIRT1 would increase chondrogenic ECM expression, as shown above for SIRT1 chemical activation. To this end we transfected TC28a2 cells with a Dox-inducible SIRT1 overexpression construct and cultured the cells in 3D pellets with or without SRT1720 (Figure 5A). Stimulation of the TC28a2 pellets with 100 nM Dox was sufficient to elicit a large increase in SIRT1 protein levels (Figure 5B). However, SIRT1 overexpression without SIRT1 activation did not increase chondrogenic gene expression (Figure 5C). Simultaneous SIRT1 overexpression and activation generated increases in transcripts for the chondrogenic genes, almost identical to those seen with SRT1720 activation alone (Figure 2 and 5C). This suggests that SIRT1 protein abundance is not the rate limiting factor in inducing chondrogenic gene expression. Rather, additional factors, acting in concert with SIRT1, contribute to its activity and enhancement of a chondrogenic phenotype in a 3D setting.

**Figure 5.**
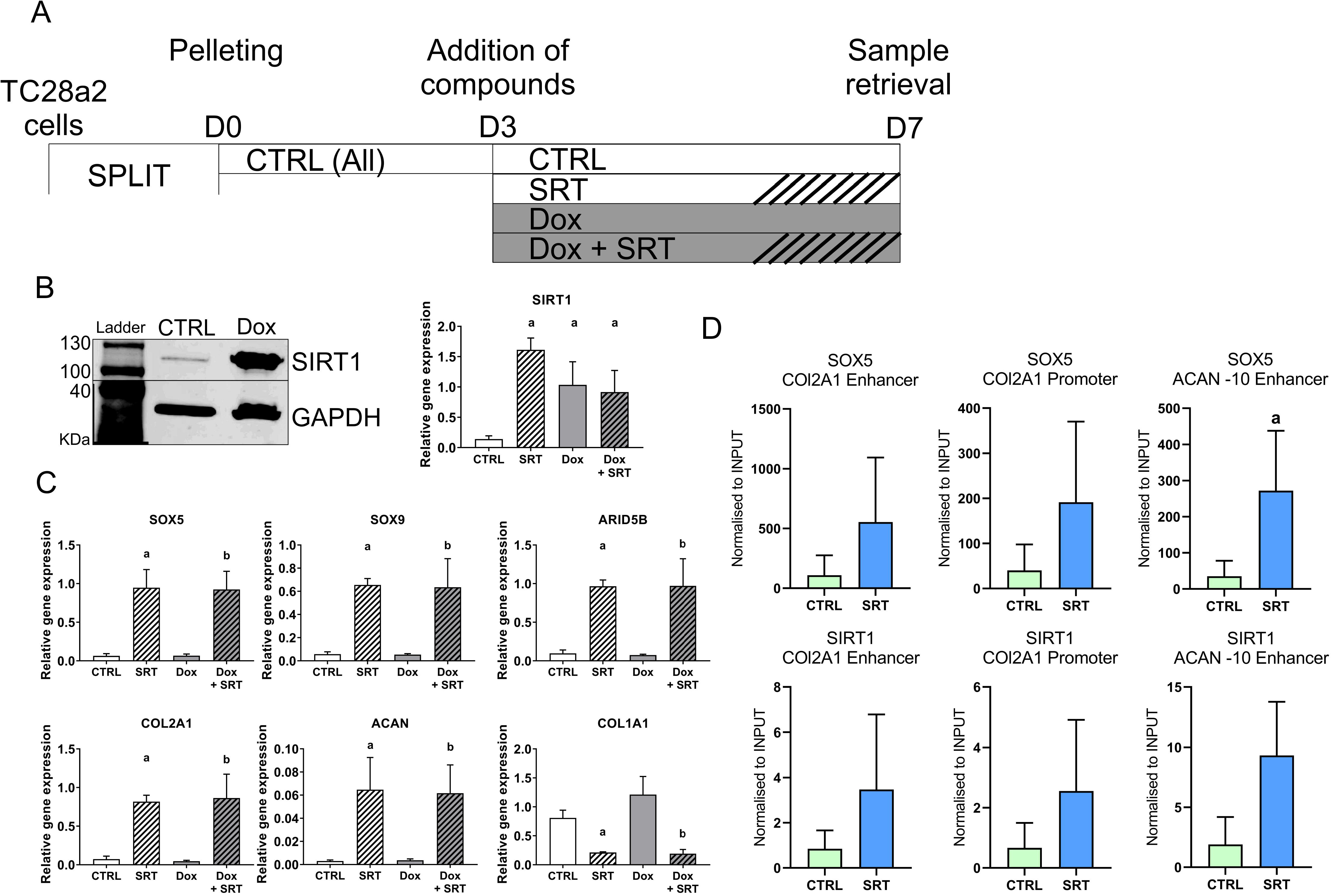
SIRT1 activation is more important than presence in promotion of chondrogenic genes. A) Schematic diagram of TC28a2 pellet culture and conditions. Bar designs for each condition are conserved between panels A, B and C. B) Western blot analysis of SIRT1 protein expression in Dox responsive SIRT1 overexpressing TC28a2 cells, treated with or without 100 μg/ml Dox from day 3 till day 14. C) QRT-PCR gene expression analysis of chondrogenic genes in Dox responsive SIRT1 overexpressing TC28a2 pellets culture for 14 days (N=4 biological repeat). Pellets were treated from day 3 till day 14 with DMSO control, SRT1720 (5 μg), Dox (100 μg/ml) or a combination of Dox and SRT1720. Data displayed relative to housekeeping gene *GAPDH*. D) ChIP analysis of SIRT1 and SOX5 occupancy of the *COL2A1* enhancer and promoter region, and of the *ACAN* −10 enhancer region in day 14+14 pellets derived from hPSC-chondroprogenitors (N=3 biological repeat). Data displayed as normalised to raw lysed sample Input. *a* indicates significance difference (p≤0.05) to DMSO control. *b* indicates significant difference (p≤0.05) to Dox treated control.

### SIRT1 activation leads to SOX5 enrichment at the *ACAN* -10 enhancer site

As SOX5 was significantly increased during SIRT1 activation in pellet culture, we further assessed the dynamics between SIRT1 and SOX5 in regulating *ACAN* expression. To this end, we carried out ChIP analysis for SIRT1 and SOX5 in day 14+14 chondroprogenitor pellets with or without stimulation with SRT1720. PCR analysis of the ChIPed DNA revealed significant enrichment of SOX5 at the *ACAN* −10 enhancer region, with SIRT1 showing a trend towards enrichment (p=0.06) following SRT1720 activation (Figure 5D). The data suggest that SIRT1 contributes to *ACAN* expression by enrichment of SOX5 on the enhancer site of the gene.

### ARID5B is required for type-II collagen gene expression in SIRT1 activated chondrogenic pellets

Importantly, the gene expression of *ARID5B* was significantly increased in SIRT1 activated pellets (Figure 2B) alongside SOX5. Thus, we investigated whether it was likely to be involved in the increased levels of aggrecan and type-II collagen observed in the developing 3D hESC-cartilage pellet culture. Pearson’s correlation analysis between *COL2A1* and *SOX9,* or *ARID5B* exhibited a weak correlation in control 3D culture. However, after SIRT1 activation we observed a significant correlation (r^2^= 0.994, p=0.0002) between *COL2A1 and ARID5B* (Figure 6A).

**Figure 6.**
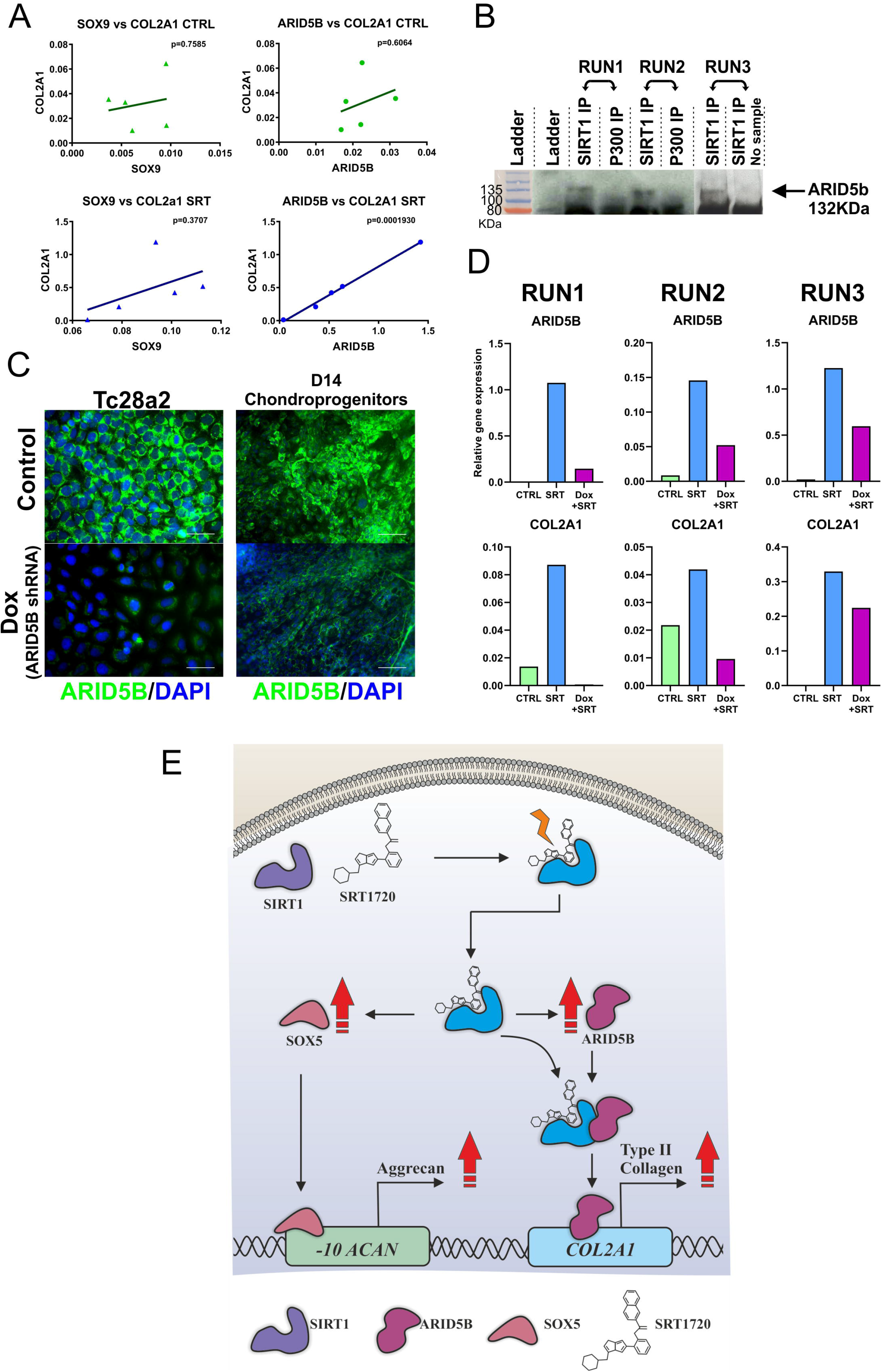
ARID5B is required for *COL2A1* expression in SIRT1 activated pellets. A) Regression curve and Pearson’s r correlation p value analysis of chondrogenic transcription factors, *SOX9* or *ARID5B* vs *COL2A1* gene expression (N=5 biological repeat) in day 14+14 pellets culture with DMSO control or SRT1720 from day 14+3 to 14+14. B) Western blot analysis of ARID5B in SIRT1 or P300 immunoprecipitate from day 14+14 pellets treated with either DMSO control or SRT1720 from day 14+3 till day 14+14. C) Immunocytochemistry of ARID5B in Dox responsive *ARID5B*-shRNA TC28a2 cells and day 14 hPSC derived chondroprogenitors, treated with DMSO control or 100ng/ml Dox for 48 hours. D) Gene expression analysis of *ARID5B* and *COL2A1* in Dox responsive *ARID5B*-shRNA hPSC derived chondroprogenitor pellets cultured for 14 days in 3D, treated with DMSO control, SRT1720 (5 μM), or SRT1720 (5 μM) and Dox (100 ng/ml) from day 14+3 till 14+14. Gene expression data displayed as relative to housekeeping gene *GAPDH.* E) Models for SIRT1 activation, including impact on downstream associated factors and resulting ECM component expression.

To assess any direct binding between ARID5B and SIRT1 we employed co-IP, with P300 used as a non-ARID5B-binding partner control. Western blot analysis of co-IP samples probed with the ARID5B antibody, displayed a band at approximately 135KDa (ARID5B size; 132KDa), detectable only in the SIRT1 immunoprecipitated lanes (Figure 6B). This band was not present in the P300 control immunoprecipitation, nor in the SIRT1 antibody only control lane.

To support this correlation of SIRT1 activation with ARID5B function, a Dox inducible *ARID5B*-shRNA construct was utilised. Upon Doxycycline stimulation ARID5B protein levels were suppressed in both TC28a2 cells and day-14 hESC-chondroprogenitors (Figure 6C). When stimulated with SRT1720 and Doxycycline, the resultant decrease in ARID5B, was reflected in a subsequent reduction in *COL2A1* expression (Figure 6D) not seen in controls. These data indicate that SIRT1 stimulates COL2A1 expression in an ARID5B-dependent manner, possibly via a novel mechanism involving direct binding of ARID5B to SIRT1.

## Discussion

The deacetylase enzyme SIRT1 has been implicated as an important regulator of cartilage homeostasis [11,13,15]. This study aimed at understanding the involvement of this enzyme and the timing of its activity during human chondrocyte development by using human embryonic stem cells.

SIRT1 is important in the maintenance of pluripotency, interacting with multiple pluripotency associated pathways including SOX2, and P53-dependent expression of NANOG and POU5F1 [31–35]. SIRT1 inhibition in hPSCs reduced stemness and promoted differentiation [36–38]. Therefore, the notable SIRT1 expression observed in the hESCs was expected. An increase in SIRT1 during the early stages of the chondrogenic differentiation protocol (days 2-5), when mesoderm induction occurs may reflect lineage commitment [39]. However, this may not be deacetylase activity-related, as activation or inhibition of SIRT1 during this period did not promote (or impair) hPSC mesodermal differentiation in our study.

This study has substantially improved our original 2D chondrogenic differentiation protocol [3–5,40]; especially by extending into a 3D pellet culture phase. Though pellet culture of hPSC derived chondroprogenitors has been achieved previously [41–43], our study utilised a defined development system, without selection from embryoid bodies [43] or transition through an MSC-like intermediary, nor the associated hypertrophic/fibroblastic gene expression [44]. Importantly, the results show pellet culture cells have enhanced chondrogenic gene expression after 14 days, with a cartilage-like, alcian blue positive ECM emerging after 28 days.

In striking contrast to in 2D, activation of SIRT1 during the 3D pellet culture significantly increased the main chondrogenic proteins in the chondroprogenitors, whilst decreasing the fibrotic and hypertrophic factors. This finding was also replicated during activation of SIRT1 in TC28a2 cultured in 3D culture. Altered SIRT1 activity and responsiveness between 2D and 3D culture has previously been shown in primary human chondrocytes [23]. Previously it was proposed that the differential response stemmed from cell de-differentiation in 2D or redifferentiation in 3D respectively. Though no study has demonstrated this in development, we showed previously that the transition of hESC-chondroprogenitors from 2D to 3D aggregates, correlates with higher COL2A1 expression [5]. Thus, the change in responsiveness to SRT1720 in 3D indicates that 3D cell architecture and/or cell contact is an essential factor for its effect on chondrogenic expressions.

Whilst activation of SIRT1 dramatically increased several chondrogenic transcription factors and ECM components, SIRT1-inhibited samples were able to differentiate into chondrocyte-like cells and produce chondrogenic ECM proteins much like non treated controls. Crucially, this is in line with *Sirt1*^−/−^ mouse KO models [17], which do form cartilage but have an altered cartilage phenotype and express decreased levels of type-II collagen and aggrecan. This supports the idea that SIRT1 deacetylase activity is not essential for chondrogenic gene expression per se, but instead is important epigenetic regulator facilitating the synthesis of adequate matrix protein during human cartilage development. This is reflected in the elevated type-II collagen and aggrecan levels in our SIRT1 activated samples.

Here we uncovered that SIRT1 overexpression combined with activation did not increase chondrogenic gene expression beyond activation alone, suggesting adequate amounts of SIRT1 in the cells but a deficit of activation factor(s). SIRT1 activation led to significant increases in the expression of SOX9 associated co-factors, in particular SOX5 [45] and ARID5B [22], indicating their potential importance as downstream regulators of chondrogenesis. There was a significant enrichment of SOX5 (with a trend to SIRT1 enrichment) at the ACAN −10 enhancer, in line with previous observations for SOX9 [13]. This together with the demonstration of SIRT1-ARID5B association, suggests that SIRT1 is reliant on these downstream factors to regulate gene transcription as summarized in Figure 6E.

SOX5 binds to and co-directs chondrogenesis in combination with SOX9 [45,46], as part of the SOX trio required for permanent cartilage [19]. Importantly this transcription factor trio is able to bind to multiple other factors to produce enhanceosomes, including P300, itself a SIRT1 target [24]. SIRT1 has also been shown to deacetylate multiple SOX family members [31,32], including SOX9, facilitating its translocation into the nucleus, and thereby promoting chondrogenic activity [13]. During chondrogenesis SIRT1 has been shown to bind both COL2A1 and ACAN promoter/enhancer sites [23], as has SOX5, specifically at the −10 enhancer region [20]. Indeed Lefebvre and Dvir-Ginzberg surmised that there are several conserved lysine residues between the SOX family members [21] particularly in the HMG box motif (www.phosphosite.org) which would be potential targets for SIRT1 modification, indicating that SIRT1 is likely to regulate nuclear entry of SOX gene family proteins to facilitate their transcriptional activity. Here we report, that ARID5B is also expressed in hESC-chondrogenic pellets undergoing differentiation. ARID5B is reported to form a complex with PHF2, directing factors such as SOX9. In this study ARID5B expression also correlated significantly with chondrogenic ECM proteins, and these observations and inferred regulatory roles suggest further potential drug targets to manipulate human chondrogenesis and promote healthy skeletal development.

In summary this work has identified a critical role for SIRT1 in control of ECM protein expression in a hESC-chondrocyte development. It has also identified, and refined our understanding of, the role of several additional factors regulating human chondrogenic development, We have shown that, in this human model of chondrogenic development sustained SIRT 1 activation differentially increases gene expression for ECM proteins, while reducing GAG formation/retention. This regulatory effect is stage dependent and requires a 3D arrangement of developing chondroprogenitors. Additionally, we have shown that after SIRT1 activation, ARID5b and COL2a1 both increase in developing chondroprogenitors and changes in these genes are directly correlated, suggesting co-regulation. Finally, SIRT1 induction of COL2A1 expression is ARID5B-dependent, and we identify a potential novel mechanism for control of cartilage protein synthesis involving binding of ARID5B to SIRT1. Thus, SIRT1 activity is a benefit to cartilage development if the positive effects on chondrogenic proteins can be balanced by other positive modulators of GAG synthesis.

## Experimental procedures

### hPSC culture

The human embryonic stem cell lines Man-7 and Man-13 [40,47] were cultured under feeder-free conditions in mTeSR1 (StemCell Technologies, France), on 6 well tissue culture plastic (TCP) plates (Corning, UK) pre-coated with 5 μg/ml Vitronectin (N-Terminal fragment Life Technologies, USA). Cells were passaged at 80% confluency using 0.5 mM EDTA and plated with 10 nM ROCK inhibitor Y27632 (Tocris, UK) exposure for a maximum of 24 hours after a split. For experimental use, hESCs were passaged at 100,000 cells /cm^2^ into a 6 well plate pre-coated with fibronectin (Millipore, USA). Differentiation protocols were started when cells were 70-80% confluence.

### Directed differentiation protocol

Chondroprogenitors were produced by differentiating hESCs through a defined differentiation protocol (DDP) (Fig 1A). For chondrogenic differentiation hESCs were differentiated in a basal medium (DMEM:F12, 2 mM L-glutamine (Life Technologies), 1% (vol/vol) insulin, transferrin, selenium (ITS) (Sigma, UK), 1% (vol/vol) non-essential amino acids (Thermo), 2% (vol/vol) B27 (Gibco), 90 μM β-mercaptoethanol) (Gibco)) supplemented with appropriate sequential addition of growth factors as previously described [3,5]; Stage 1 (day 1-3):CHIR (2 μM, R&D systems, UK), Activin-A (reducing from 50-10 ng/ml, Peprotech, UK) and BMP2 (Concentration set at EC60 for consistent activity; R&D systems), followed by Stage 2 (day 4-8): BMP2 (See above), SB431542 (1 mM) and GDF5 (20 ng/ml) (all Peprotech) and finally Stage 3 (day 9-14): GDF5 (20 rising to 40 ng/ml at day 11), FGF2 (20 ng/ml) ml (both Peprotech), with BMP present at half stage 1 concentration until day 11. At days 4 and 8, cells were split 1 in 4 using EDTA and TrypLE respectively. Samples were collected for RNA and protein at day 4, 8 and 14 [4].

Cells cultured in the above medium were supplemented with, DMSO vehicle control (1 μl/ml), SRT1720 (Selleckchem, UK)[48] (concentration specified in figure legend) a proven SIRT1 enzyme activator, or EX527 (Sigma) [13,27] (concentration specified in figure legend), an inhibitor of SIRT1, at either days 2-5, in stage 3 (days 9-14), or during pellet culture (below) as indicated in Results.

### Pellet formation

At day 14 cells were separated from coated tissue culture plates using TrypLE (Life Technologies) and counted using a nucleocounter (Chemeotech, NL). Cells were resuspended at 5×10^5^ cells/ml in the same medium used at day 14. A 1 ml aliquot of cells was pipetted into each 15 ml centrifuge tube and centrifuged at 300 xRCF for 3 minutes to sediment cells; caps were left loose, and pellets were incubated at 37°C 5% CO_2_ for 3 days. Pellets were then transferred to chondrobasal medium (CB) containing Dulbecco’s Modified Eagle Medium (DMEM) (Life Technologies), L-glutamine (2 mM), L-Proline (40 μg/ml)(Sigma), 1x ITS (Sigma), ascorbate-2-phosphate (50 μg/ml)(Sigma), and dexamethasone (100 nM) (Sigma) supplemented with TGFβ3 (10 ng/ml)(Peprotech), GDF5 (20 ng/ml)(Peprotech) and BMP2 (half concentration of stage 1) (R&D systems). Pellets were separated into 4 groups and incubated with the above medium containing DMSO vehicle control (1 μl/ml), 5 μM SRT1720 (concentration based on results described in Supplementary figure 2A), and 5 μM EX527 as indicated in Results. Medium was changed every 3 to 4 days until either 14- or 28-days post pelleting (See Fig 2A).

### TC28a2 cell culture

Immortalised chondrocytes, TC28a2 cells [49], were cultured in DMEM containing 10% FBS, and L-glutamine (2 mM). Cells were cultured on TCP in 2D. For 3D culture, cells were trypsinzed, suspended in TC medium at 1×10^6^ cells/ml and centrifuged at 300 xRCF for 5 minutes. Pellets were incubated for 3 days then medium changed to CB media as described previously.

### Gene transcription analysis

Cells were transferred to RLT buffer directly and RNA extracted by use of a RNeasy QIAgen kit according to manufacturers’ instructions. For 3D cultures, pellets were first dissociated by grinding with Molecular Grinding Resin™ (Sigma) with a pestle in a 1.5 ml microcentrifuge tube (MCT). The ground pellets were then mixed with RLT buffer and RNA extracted. Extracted RNA was Dnase treated and RT reaction used to produce cDNA with ABI-RT kit (Life technologies). Quantitative real-time polymerase chain reaction (qRT-PCR) was conducted using PowerUp™ SYBR™ green (Life technologies) and primers for chondrogenic and non-chondrogenic genes (see Table S1). Data was calculated and displayed as relative expression to housekeeping gene *GAPDH*.

### Western blotting and immunoprecipitation

Cell samples were lysed in RIPA buffer (Sigma) with added 1x Roche cOmplete protease inhibitors (Sigma) and quantified using Pierce^®^ BCA (Thermo). Aliquots containing 30 μg of protein were boiled at 95°C for 10 minutes. Samples were loaded on 10% SDS PAGE gels (Invitrogen). Protein gels were first blocked in 5% Marvel milk™ solution in PBS-T before probing with specific primary antibodies (see Table S2). Antibody binding was detected using LICOR IRDye secondary antibodies. Blots were imaged using the Odyssey CLx. Quantification of protein was achieved using ImageJ, and protein quantification was calculated relative to GAPDH internal control levels.

For immunoprecipitation procedures, approximately 15-20 pellets were fixed for 1hr in 4% paraformaldehyde, then washed twice before being ground (as above). Material was lysed using 300 μl RIPA buffer containing PMSF (1 mM) and protease inhibitor cocktail (Roche). Samples were centrifuged at 15,000 xRCF for 10 minutes at 4°C, and supernatant assessed for protein concentration. A 100 μg aliquot was diluted to 300 μl and incubated overnight with either antibody to SIRT1 (Millipore) or to P300 (Abcam) at 4°C under constant agitation. After preliminary incubation, 50 μl of washed magnetic beads (Pierce) were added to the solution and incubated at 4°C for a further 5 hours. Supernatant was removed, and the beads washed 3 times in PBS before being boiled at 95°C for 10 minutes with reducing buffer (Thermo, 39000). Resulting samples was analysed using western blotting with detection using anti-rabbit IgG light chain antibody (Abcam) and ClarityMax ECL or DAB (BIO-RAD and Sigma respectively).

### Chromatin immunoprecipitation (ChIP)

ChIP was conducted using the Diagenode LowCell ChIP kit (Diagenode). Chondrogenic pellets were washed in PBS, then fixed in 1% PFA at room temperature for 30 minutes. After incubation, glycine was added to a final concentration of 0.125 M, the solution was mixed and further incubated at room temperature for 5 minutes. Samples were then washed twice in ice cold PBS containing inhibitor cocktail, PMSF (1 mM supplier) and protease inhibitors (supplied by kit supplier), Sodium Butyrate (5 mM). Pellets were then ground using pestle and Molecular Grinding Resin (as above) and resuspended in 60 μL buffer B supplied in kit. Samples were sonicated using the BioRupter (Diagenode) at full power for 30 cycles (30 seconds on; 30 seconds off). Chromatin was extracted using SIRT1 and SOX5 antibodies (see Table S2), and DNA isolated as described in kit. Isolated DNA was analysed using qRT-PCR using primers for *COL2A1* promoter and enhancer, and *ACAN* −10 enhancer (see Table S1).

### Lentiviral overexpression

Stably expressing TC28a2 and Man-13 cells were generated using an established lentiviral method (SBI Systems Bioscience). cDNA sequences used for lentiviral overexpression were based on that of human *SIRT1* (Gene ID: 23411) and were subcloned into a modified Doxycycline (Dox) inducible viral expression vector based on the pCDH-EF1-T2A-copGFP vector. His-Flag-SIRT1 (gift from Prof. Danny Reinberg, NYU, NY) and *ARID5B*-shRNA expression was controlled by the inducible TRES3G promoter. Tetracycline (Tet)-On 3G trans-activator protein and tagGFP2 fluorescent protein were under the constitutive promoter Elongation factor 1 (EF1a). This system is responsive to Tet and derivative Dox. HEK293T cells were co-transfected with psPax2, pMD2.G packaging as well as a target pCDH vector. Production of virus particles was induced by addition of 10 mM sodium butyrate (Millipore) for 4 hours, 24 h post-transfection, and virus particles were harvested from the medium after 48 hours and filtered through a 0.2 μm filter prior to addition to target cells. TC28a2 or Man-13 cells were then infected with virus using 5 μg/ml protamine sulphate (Sigma). Cells were passaged, followed by fluorescence-activated cell sorting (FACSAria Fusion, Beckon-Dickenson).

### Dimethyl-methylene blue (DMMB) sulphated GAG assay

DMMB assay was undertaken using the Blyscan™ assay kit (Biolcolor), to ascertain the sulphated glycosaminoglycan (GAG)-content of pellets. Pellets were first digested in 200 μl papain digestion solution containing PBS and 0.5 mg/ml papain (Sigma) overnight at 65°C with intermittent agitation. Samples were centrifuged at 10,000 xRCF for 10 minutes and supernatant decanted for storage before use. Digested samples were stored at −20°C until assayed.

For assay, 50 μl of sample was diluted with 500 μl of Blyscan dye reagent and incubated for 30 minutes. Samples were centrifuged at 12,000 xRCF for 10 minutes and resulting pellet resuspended in 500 μl dissociation reagent. 100 μl of samples was measured for absorption at 650 nm. A Quant-iT pico-green dsDNA assay (Thermo) was used to quantify DNA, which was used to standardise readings.

### Histological assessment (IHC, IP)

Chondrogenic pellets were harvested at Day 14 monolayer culture, and at −14 days or −28 days in pellet culture, rinsed in PBS and fixed overnight in 4% paraformaldehyde at 4°C. After which, pellets were stored in 70% ethanol before processing. Processed samples were embedded in PPFA wax, sectioned to 5 μM, and deposited on glass slides.

Wax was removed by Xylene and sections were rehydrated through serial alcohols (100, 90, 70%) then dH_2_O. Sections were washed 3 times in PBS- 0.1%Tween 20 then subject to antigen retrieval with citrate buffer (pH6.5) and heated to 95°C, or Pepsin enzyme (Thermo). Sections were first blocked in primary animal serum, then incubated overnight at 4°C with primary antibodies as described in results (see Table S2), with antibody detection by either biotinylated secondary followed by streptavidin peroxidase or fluorochrome tagged secondary antibody as appropriate. No primary, or inappropriate primary antibodies were used as control. A hyaluronic acid binding protein (HABP) antibody was used to visualise the polysaccharide hyaluronic acid.

### Statistical analysis

All statistical analysis was run using Prism Graph-pad. Gene and protein expression changes were analysed using Mann-Whitney U test. ChIP was analysed using a ratio paired t-test. Correlation analysis was quantified using a Pearson’s correlation coefficient. A p-value of ≤ 0.05 was considered as statistically significant.

## Acknowledgments

This work was supported mainly by Arthritis Research UK (Grants R20786) to SJK and MDG, an MRC UKRMP hub award (Grant MR/K026666) to SJK and a Rosetrees Trust grant A1984 to CAS, SJK and MDG. We thank Mr. Peter Walker for histology training Dr Ashok Kumar for ChIP training. TC28a2 cells were kindly donated by Dr Louise Reynard of Newcastle University. Aggrecan G1 antibody was generously donated by Prof Tim Hardingham of University of Manchester.

## Author Contributions

Experimental design and undertaking by CAS. Cell culture by CAS and NB. Additional samples supplied by PH and MN. Virus design and production by SC. Study design and concept by MDG and SJK. Manuscript written and edited by CAS, MDG and SJK.

---

**Figure S1: Influence of SIRT1 modulators on early hESC mesoderm and chondroprogenitor differentiation during 2D culture**. A) QRT-PCR gene transcription analysis of alternate epigenetic factors associated with initial hESC-chondrogenesis in the 14-day DDP (N=4 biological repeat). B) Western blot protein expression analysis of PHF2 and GAPDH for cell samples taken at days 0, 4, 8 and 14 during the chondroprogenitor differentiation protocol. C) QRT-PCR analysis of samples taken for hESCs, end of stage 1 (day 4), end of stage 2 (day 8) and end of stage 3 (day 14) of the 2D chondroprogenitor differentiation protocol for cells treated with the selective SIRT1 activator, SRT1720, or inhibitor, EX527 between days 2-5 of the chondrogenic protocol (N=4 biological repeat), and D) Western blot protein expression analysis of SIRT1, SOX9, SETD7, PHF2 and GAPDH for cell samples taken at day 0, 4, 8 and 14 for cells treated with 5 μM EX527 during days 2-5 of the protocol (N=1). E) QRT-PCR analysis of samples taken at the end of stage 3 (day 14) of the 2D chondroprogenitor differentiation protocol for control cells or cells treated with selective SIRT1 activator SRT1720 or inhibitor EX527 from day 8 onwards (N=4 biological repeat). All gene expression data displayed relative to housekeeping gene *GAPDH.* * signifies significant difference (p≤0.05) compared to day 0.

**Figure S2: Effect of epigenetic modulators in 3D pellet culture**. A) Gene expression analysis of day 14+7 chondrogenic pellets treated with 1 or 5 μM SRT1720 for 4 days from day 14+3 (N=3 biological repeat). B). QRT-PCR gene expression analysis of chondrogenic transcription factors in pellets treated with DMSO control, or SRT1720 from days 3-14 (11 days) or from days 11-14 (3 days) (N=5 biological repeat). SRT1720 concentrations as displayed. C) QRT-PCR gene expression analysis of day 14+14 pellets treated with DMSO control (CTRL), SIRT1 activator SRT1720 (5 μM), or SETD7 inhibitor (r)-PFI-2 (1 μM) from days 14+3 till 14+14 (N=1). Gene expression data displayed as relative to housekeeping gene *GAPDH. a* signifies significant difference (p≤0.05) compared to DMSO control (CTRL).

**Figure S3. Activation of SIRT1 changes balance in expression of ECM components**. A) DMMB GAG concentration assay for D14+28 pellets treated with DMSO and SIRT1 activator SRT1720. B) QRT-PCR gene expression analysis of HAS2 in day 14 2D pre-pellet cells, and pellets at day 14+14, treated with DMSO control or SIRT1 activity modulators from day 14+3 to 14+14 (N=7 biological repeat). Data displayed as gene expression relative to housekeeping gene *GAPDH* and shown as box plots. *a* indicates significant difference to Day 14 pre-pellet sample (p≤0.05), *b* indicates significant difference to day 14+14 DMSO vehicle control (p≤0.05). C) IHC and IF pellet staining for type-II collagen, HAS2 (example positive cells highlighted with arrows), HABP and lubricin in day 14+28 pellets in 3D cultures treated +/− SRT1720 (5 μM).

