## Supplementary figures and images for "SIRT1 activity orchestrates ECM expression during hESC-chondrogenic differentiation"

### Figure S1

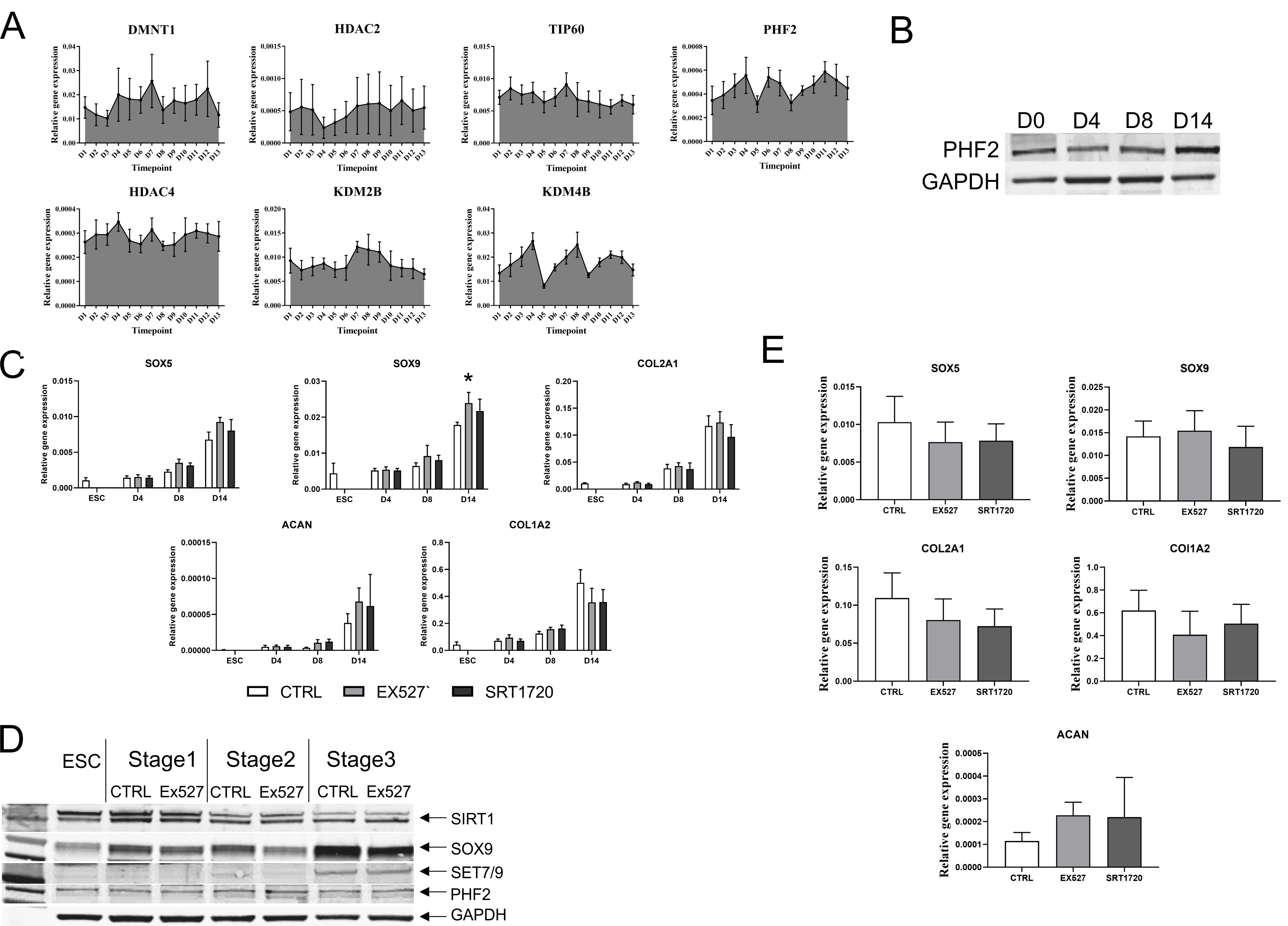

### Figure S2

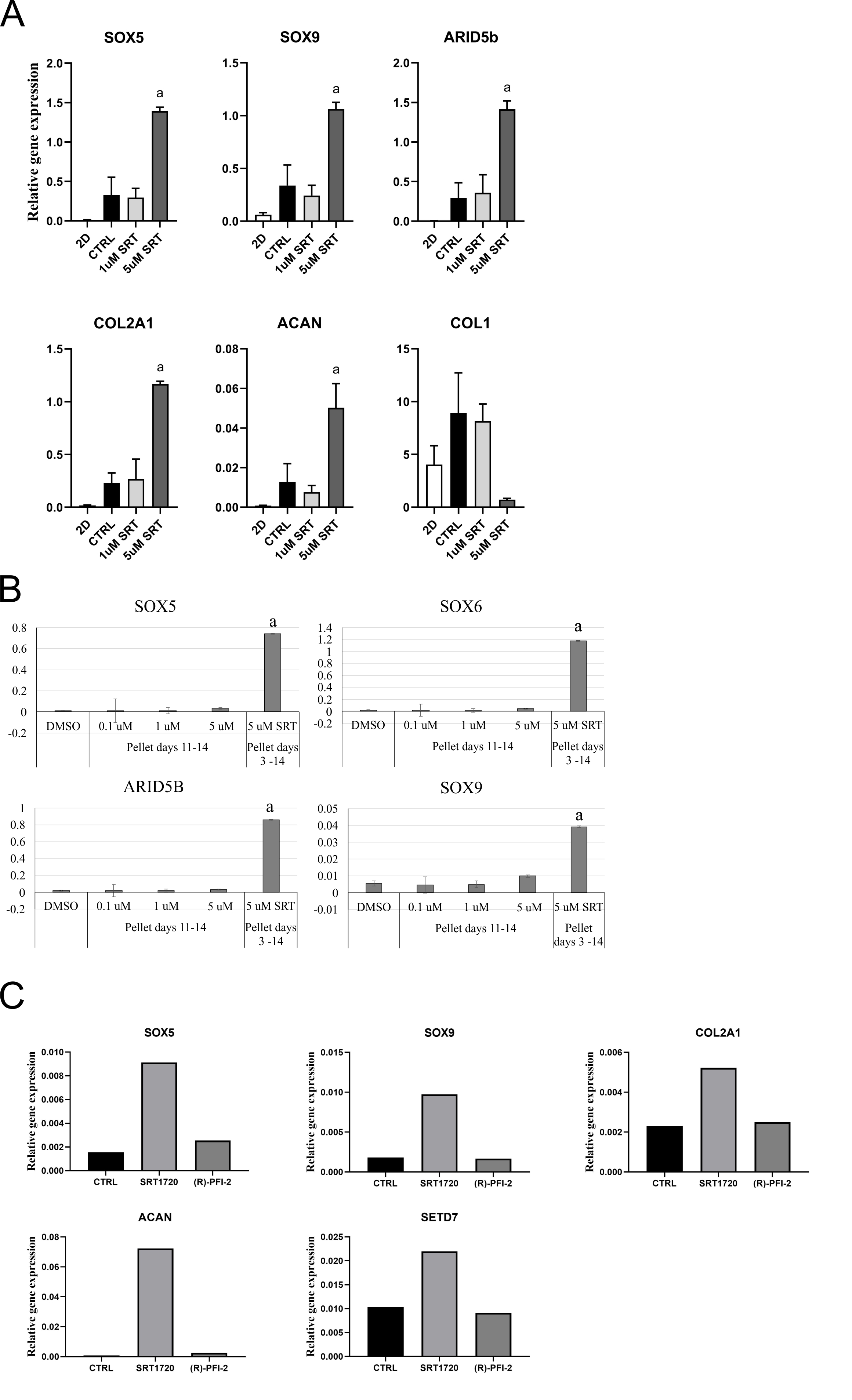

### Figure S3

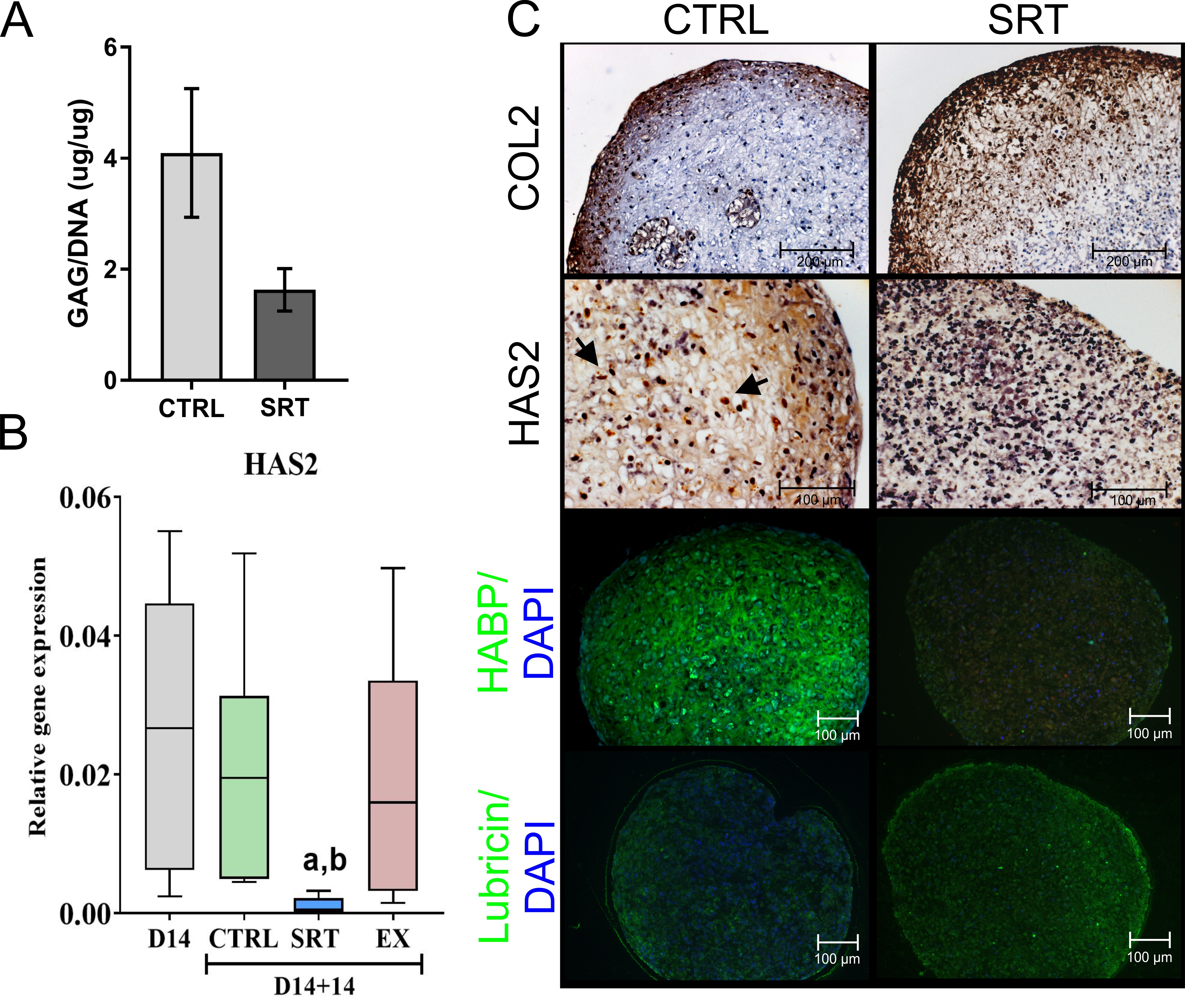
