## Supplementary material for "SIRT1 activity orchestrates ECM expression during hESC-chondrogenic differentiation": Table 1, Table 2

| GENE | FORWARD | REVERSE |
| --- | --- | --- |
| ACAN | CATTCACCAGTGAGGACCTCG | TCACACTGCTCATAGCCTGCTTC |
| ARID5A | CCTCTGGAAGAACGTGTACGA | CTTCAGGTGCCGCACGTAT |
| ARID5B | GAGCAAAGGCATCTCCCAGT | GTTGAGGCCCGAGTTACACA |
| COL1A2 | CAGCCGCTTCACCTACAGC | TTTTGTATTCAATCACTGTCTTGCC |
| COL2A1 | GGCAATAGCAGGTTCACGTACA | CGATAACAGTCTTGCCCCACTT |
| DNMT1 | GATCGAATTCATGCCGGCGCGTA  CCGCCCCAG | ATGGTGGTTTGCCTGGTGC |
| GAPDH | ATGGGGAAGGTGAAGGTCG | TAAAAGCAGCCCTGGTGACC |
| HDAC2 | CGGTGTTTGATGGACTCTTTG | CCTGATGCTTCTGACTTCTTG |
| HDAC4 | GAGAGACTCACCCTTCCCG | CCGGTCTGCACCAACCAAG |
| KDM2B | GTTAGTGGTAGTGGTGTTTTGG | AGCAGATGTGGTGTGTGGTC |
| KDM4B | CGCGGCAGACGTATGATGAC | TGTCATGGCCTTCTTCTGGA |
| NANOG | GGCTCTGTTTTGCTATATCCCCTAA | CATTACGATGCAGCAAATACAAGA |
| PHF2 | CTAAGTTGTCCCAGCAGGAGG | GTTTGGAGACGATGCGGAGG |
| POU5F1 | AGCGAACCAGTATCGAGAACC | CTGATCTGCTGCAGTGTGGGT |
| SETD7 | CGTGGTGTGCCTGAGCCC | TGAAGGAGTGATTTGCCTTGT |
| SIRT1 | GCTTATTTGTCAGAGTTCCCACCC | CAGCATTTTCTCACTGTTCCAGCC |
| SIRT2 | GAACGCTGTCGCAGAGTCATC | GGTTGGCTTGAACTGCCCAG |
| SIRT3 | CAGTCTGCCAAAGACCCTTC | CAACCACATGCAGCAAGAAC |
| SIRT6 | TTGTGGAAGAATGTGCCAAG | CCTTAGCCACGGTGCAGAG |
| SIRT7 | TCACCCACATGAGCATCACC | GGAACGCAGGAGGTACAGAC |
| SOX5 | ATCCCAACTACCATGGCAGCT | TGCAGTTGGAGTGGGCCTA |
| SOX9 | GACTTCCGCCACGTGGAC | GTTGGGCGGCAGGTACTG |
| TIP60 | CAGGACAGCTCTGATGGAATAC | AGAGGACAGGCAATGTGGTGAG |
| SOX6 | GCAGTGATCAACATGTGGCCT | CGCTGTCCCAGTCAGCATCT |
| RUNX2 | GACGAGGCAAGAGTTTCACC | GCCTGGGGTCTGTAATCTGA |
| COLX | CACCAGGCATTCCAGGATTCC | AGGTTTGTTGGTCTGATAGCTC |
| COL2A1-PROMOTER | CGCTGGGCTGTAACCTGAAC | GGAAGCGTGACTCCCAGAGA |
| COL2A1-ENHANCER | ATCCTCCTTTGTGAGGCTTGTT | AGTACGAGAGAACCCACTGGAC |
| ACAN-ENHANCER | ATGTGTCTCAAGTCCAGAATGGAA | GAAATTCCTTTAGCGGCAACGCCT |

**TABLE 1 – List of DNA primer sequences**

| TARGET | PRODUCT CODE | COMPANY | HOST |
| --- | --- | --- | --- |
| SIRT1 | 07-131 | Millipore | Rb |
| SETD7 | ab14820 | Abcam | M |
| PHF2 | ab154983 | Abcam | M |
| SOX9 | ab5535 | Millipore | Rb |
| GAPDH | #5174 | Cell Signal | Rb |
| ARID5B | HPA015037 | Sigma | Rb |
| KAT3B/P300 | ab14982 | Abcam | M |
| SOX5 | ab94396 | Abcam | Rb |
| HAS2 | ab140671 | Abcam | M |
| TYPE II COLLAGEN | ab185430 | Abcam | M |
| LUBRICIN | ab28484 | Abcam | Rb |
| GRC5/PHF2 | ab65771 | Abcam | Rb |
| HABP | ab53842 | Abcam | Sh |
| Aggrecan G1 domain | [27] | in house | Rb |
| Anti-Rabbit IgG light chain (HRP) | ab99697 | Abcam | M |

**TABLE 2 – List of Antibodies**
